# Tracking mobilization uncovers an evolutionarily conserved mechanism in suppressing transpositions during somatic development

**DOI:** 10.1101/2025.02.07.636160

**Authors:** Yi Ni Luo, Yu Liang, Shuheng Wu, Yazi Deng, Wei Wu, ZZ Zhao Zhang, Wei Wu, Lu Wang

**Affiliations:** Key Laboratory of RNA Innovation, Science and Engineering, Shanghai Institute of Biochemistry and Cell Biology, Center for Excellence in Molecular Cell Science, Chinese Academy of Sciences, University of Chinese Academy of Sciences, Shanghai, 200031, China; Key Laboratory of Multi-Cell Systems, Shanghai Institute of Biochemistry and Cell Biology, Center for Excellence in Molecular Cell Science, Chinese Academy of Sciences, University of Chinese Academy of Sciences, Shanghai, 200031, China; Department of Pharmacology and Cancer Biology, Duke University School of Medicine, Durham, NC, USA; Duke Regeneration Center, Duke University School of Medicine, Durham, NC, USA

## Abstract

Transposon mobilizations could cause gene mutations and even genomic instability, whose aberrant activation is tightly linked with cancer, neurodegenerative disorder and other pathologies. However, the precise suppression mechanisms of transposon mobilizations during somatic development throughout evolution remain largely obscure. Here, by spatiotemporally monitoring transpositions with single-cell resolution, we determined a highly conserved mechanism in suppressing transposon mobilizations. We identified that Cramp1 safeguards the genomic integrity in both *Drosophila* hindgut regeneration and mouse embryonic erythropoiesis through silencing transposon mobility. Cramp1 is characterized as a major hallmark that specifically binds to the DNA sequences of histone gene cluster to initiate transcription of linker histone H1, which subsequently promotes H3K36me2-established heterochromatin for transposon silencing. Our finding highlights that with the endless arms race between hosts and transposons during evolution, the core mechanisms will also evolve to suppress transposon mobilizations during somatic development.

**One-sentence abstract:** Cramp1 plays an evolutionarily conserved role in specifically initiating linker histone H1 transcription and triggering H1-H3K36me2-established heterochromatin to suppress transposon mobilizations and maintain genome integrity during somatic development.

## Main text

Transposons have thrived in the genomes of nearly all eukaryotic organisms, occupying a whopping 46% of the human genomic DNA(*1, 2*). During evolution, domesticated transposons provide tremendous sources to rewire gene regulatory networks, shape 3D chromatin architecture and facilitate genome innovation(*3–8*). Nevertheless, being capable of replicating and mobilizing, transposon activation and insertions have long been considered as detrimental to the hosts, by driving aging and causing genetic disease, neurodegenerative disorder and cancer(*9–15*). Notably, while LINE-1 could substantially increase its mobilization rate during colorectal tumorigenesis, the LINE-1 retrotransposition events can be frequently detected in normal colorectal epithelium, contributing to somatic mosaicism(*16*).

In somatic cells, the ectopic transcription of transposons is precisely controlled by an epigenetically stable and heritably repressive structure, known as heterochromatin. H3K9 trimethylation (H3K9me3), a crucial mark associated with constitutive heterochromatin, confers the core function in maintaining this repressive structure for transposon silencing, thereby ensuring genome stability(*17–19*). In addition to H3K9me3 mediated transposon suppression, H3K36me2 and H3K27me3 were also demonstrated to repress transposon activity in other circumstances(*20–23*). While the epigenetic state is primarily maintained and dynamically regulated by the post-translational modifications of the core histones, recent evidences suggest that the linker histone H1 also contributes to epigenetic regulation of transposon activity(*24–30*).

Despite the knowledge of heterochromatin in suppressing transposon expression, lacking a genetic tool to trace retrotransposition has dampened our understanding of the mechanisms in suppressing mobilizations during somatic development. By leveraging a transposition reporter, we sought to characterize how retrotransposon mobilizations were spatiotemporally suppressed during animal development.

## Results

### dCramp1 suppresses transposon mobilizations during *Drosophila* ileum regeneration

As selfish elements, the ultimate purpose of retrotransposons is to accomplish their propagation, by inserting new copies of themselves into host genomic DNA(*15, 31*). During *Drosophila* oogenesis, *HMS-Beagle* was identified as the most active retrotransposon at mobilization level upon piRNA pathway perturbation(*32, 33*). Nevertheless, whether *HMS-Beagle* is capable of amplifying itself in the genome during somatic development remains elusive. Inspired by our previous studies that employ an eGFP reporter to monitor retrotransposon mobilizations(*32, 34*), we engineered a transposition reporter for *HMS-Beagle* (Fig. 1A). Within the cells harboring this transposition reporter, the eGFP can only be produced when the intron-spliced *HMS-Beagle* mRNA generates a new DNA copy in the genome (Fig. 1A).

**Fig.1.**
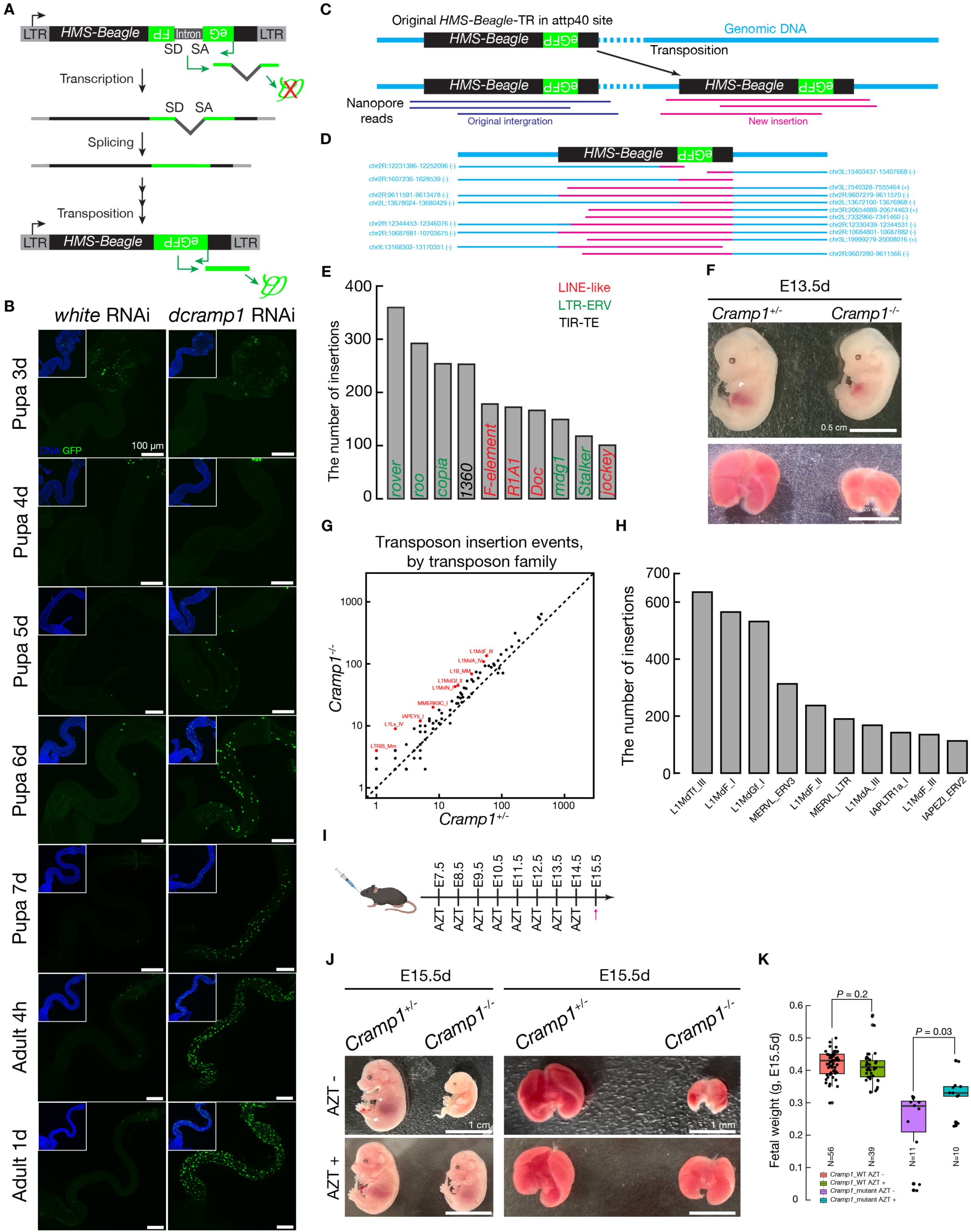
dCramp1/CRAMP1 suppresses transposon mobilizations during somatic development. A, Schematic design of an eGFP reporter to track *HMS-Beagle* transposition. The eGFP reporter is inserted in the 3’ end of *HMS-Beale* in antisense orientation. The eGFP sequence contains a disruptive intron, which is in the same direction as *HMS-Beagle* but opposite direction to eGFP. eGFP protein can only be produced upon transposon transcription, intron splicing and reverse transcribed cDNA integrating back into the genome. SA, splicing acceptor; SD, splicing donor. B, Monitoring *HMS-Beagle* mobilization events from 3-day-old pupae to 1-day-old adults in *white* and *dcramp1* depleted *Drosophila* ilea. Each GFP positive cell indicates *HMS-Beagle* mobilized at least once in the genome. RNAi was driven by *byn*-Gal4 and pupae were kept at 18 °C. C, The workflow to classify the integration events from engineered *HMS-Beagle* transposition reporter. The reads from Nanopore sequencing were identified as novel mobilizations if they contain flanking sequences mapped to the genome. D, The new insertions detected from *HMS-Beagle* transposition reporter in *dcramp1* depleted hindguts. Reads contain flanking sequences from non-repetitive genomic regions at one end or both ends of *HMS-Beagle*. E, The new insertions detected from the top 10 most active transposon families in *Drosophila* hindguts upon *dcramp1* depletion. F, Appearance of *Cramp1*^+/-^ and *Cramp1*^-/-^ embryos and fetal livers at E13.5d. G, Scatter plot showing the transposon insertion events from E13.5d *Cramp1*^+/-^ and *Cramp1*^-/-^ fetal livers. Red dots: The top transposon families increased their mobilization rate in *Cramp1*^-/-^ comparing to *Cramp1*^+/-^. *Cramp1*^+/-^ and *Cramp1*^-/-^ are siblings. H, The new insertions detected from the top 10 most active transposon families in E13.5d *Cramp1*^-/-^ fetal livers. I, Workflow showing AZT treatment of pregnant female mice. J, Appearance of *Cramp1*^+/-^ and *Cramp1*^-/-^ embryos and livers at E15.5d with or without AZT treatment. K, Box plots showing the fetal weight of wild type and *Cramp1* mutant embryos at E15.5d with or without AZT treatment. *Cramp1*_WT: *Cramp1*^+/+^ and *Cramp1*^+/-^; *Cramp1*_mut: *Cramp1*^-/-^. The central lines represent median values, the box edges represent minimum and maximum values and the whiskers show the interquartile range of the data. Two-tailed *t*-tests were used to evaluate the difference between AZT treated and untreated groups.

When tracking mobilizations, we found that only several GFP positive cells can be visualized during *Drosophila* ileum regeneration (Fig. 1B), indicating *HMS-Beagle* activity is predominantly suppressed. While such low jumping events were detected from *HMS-Beagle*, it provides a great opportunity to identify the factors that control transpositions during somatic development. After performing an RNAi screening in *Drosophila* hindgut, we found a suppressor, dCramp1 (Crm), can sufficiently prevent *HMS-Beagle* mobilizations during ileum regeneration (Fig. 1B and fig. S1A). Intriguingly, the function of dCramp1 in suppressing *HMS-Beagle* transpositions is spatiotemporally dependent. Even without dCramp1 protection, *HMS-Beagle* did not jump at the early stages of ileum regeneration (Fig. 1B and fig. S1A, pupa 3d and 4d). *HMS-Beagle* started mobilizing at pupa 5d and reached to the highest mobilization activity in the ilea of newly eclosed flies upon *dcramp1* depletion (Fig. 1B and fig. S1A). dCramp1 was classified as a *Polycomb*-group protein and the overexpressed dCramp1 exhibited a nuclear localization pattern in *Drosophila* salivary gland(*35–37*). Similarly, by inserting a GFP tag into the C-terminal of endogenous dCramp1, we found that dCramp1 exclusively localized in the nucleus and exhibited a prominent single bright locus of labeling (fig. S1, B and C). The function of dCramp1 in suppressing *HMS-Beagle* mobilizations was further validated by utilizing two different RNAi alleles and rescue experiment (fig. S1, D and E).

To capture the bona fide mobilization events from our *HMS-Beagle* transposition reporter within *Drosophila* hindguts, we employed Nanopore sequencing technology (Fig. 1C). Upon *dcramp1* depletion, we detected 13 potentially new insertions from *HMS-Beagle* reporter in adult hindguts (Fig. 1D and fig. S2, A and B). As expected, in the ovaries, with transposons being silenced by piRNA pathway, only one *HMS-Beagle* insertion was detected (fig. S2A). By further mining our Nanopore sequencing data, we characterized 4,244 novel insertions that specifically occurred in hindguts upon *dcramp1* depletion. Notably, the top 10 transposon families comprised 47.78% (2,028 insertions) of the total transposition events (Fig. 1E and fig. S2C). Futhermore, we found that those integration events were almost evenly distributed across the genome, with little differences among main chromosomes (fig. S2, D and E). In cultured-cell retrotransposition assays, researchers have discovered that LINE-1 and Alu elements could generate target site duplications (TSDs) upon integrating into genomic DNA(*38, 39*). When analyzing the spanning reads from Nanopore sequencing data, we found that only 8.26% (18/218) of the total spanning reads support retrotransposon mobilizations generating TSDs in *Drosophila* hindguts (fig. S2F). However, 89.9% (196/218) of the total spanning reads support transposon mobilizations causing genomic depletions in *Drosophila* hindguts (fig. S2F). Altogether, we concluded that dCramp1 is essential for suppressing transposon mobilizations during the *Drosophila* ileum regeneration process.

### CRAMP1 controls transposon mobilizations in mouse fetal liver

dCramp1 contains a DNA binding domain, SANT domain, which is highly conserved among fly, mouse and human (fig. S3A). To demonstrate whether CRAMP1 also suppresses transposon mobilizations during mouse somatic development, we mutated *Cramp1* in the genome (fig. S3B). None of the survival *Cramp1*^-/-^ pups were ever gotten, indicating that *Cramp1* homozygous mutants were embryonically lethal (fig. S3C). After dissecting pregnant mice at different time points, we observed that most of the fetuses could survive until E13.5d, but died at around E15.5d (Fig. 1, F and J and fig. S3D).

From those *Cramp1*^-/-^ fetuses, the only one overt phenotype we detected was anemia (Fig. 1, F and J and fig. S3D), most likely caused by the defects of erythropoiesis in fetal liver(*40, 41*). To assess whether CRAMP1 suppresses transposon mobilizations, we extracted the genomic DNA from the fetal liver and performed Nanopore sequencing. In wild type conditions, we already could detect transposon mobilizations, indicating transposons contribute to somatic mosaicism in mouse fetal liver (Fig. 1G). Comparing to *Cramp1*^+/-^ group, we detected much more transposon insertions in *Cramp1*^-/-^ fetal liver at E13.5d (Fig. 1G). Notably, all of the most active transposons were retrotransposons and more than half of them were LINE-1 families (Fig. 1H), composing nearly 20% of the human and mouse genomes(*9*).

Assuming anemia of *Cramp1* mutated fetuses was caused by retrotransposon mobilizations, then blocking reverse transcription would be expected to rescue this defect. To support this hypothesis, we treated the pregnant mice with azidothymidine (AZT) (Fig. 1I), an inhibitor of reverse transcriptase, to prevent LINE-1 transpositions(*42*). Notably, comparing to the group without AZT treatment, feeding the pregnant mice with AZT definitely rescued the severe anemia phenotype (Fig. 1J) and significantly improved the weight of *Cramp1* mutated fetuses (Fig. 1K). Additionally, we observed 9 out of 10 fetuses (90%) contained livers from AZT treated *Cramp1* mutants at E15.5d, but only 4 out of 11 (36.4%) fetuses had livers without AZT treatment. These data suggest that retrotransposon mobilizations in *Cramp1* mutated fetal liver contribute to the anemia phenotype.

### dCramp1/CRAMP1 protects genome integrity from transposon mobilizations

We previously have suggested that transposon mobilizations in germline stem cells and oocytes induced considerable DNA breaks(*32, 43*). In *Drosophila* hindguts, our data indicated that majority of the transpositions caused genomic deletions (fig. S2F), which would potentially lead to DNA breaks. Indeed, we found that DNA breaks were significantly increased upon transposon mobilizations in *dcramp1* depleted ilea, as evidenced by strong signals of TUNEL and γ-H2Av staining (fig. S4, A and B). Clear signals of γ-H2Av could be detected in the *dcramp1* depleted ilea at pupa 4d and 5d, but not pupa 3d, further indicating transposons did not propagate at the early stage during ileum regeneration (fig. S4A and Fig.1B).

To further examine whether transposon mobilizations in *Cramp1* mutants will also cause DNA breaks, we performed γ-H2AX staining in E13.5d fetal liver. Strong signals of γ-H2AX could be clearly detected in *Cramp1* mutated fetal liver (fig. S4C). However, we did not observe any γ-H2AX signals in fetal kidney and heart (fig. S4C), suggesting CRAMP1 probably specifically functions to suppress transposon activity in fetal liver.

### dCramp1 silences transposon transcription in fly hindgut

One possible role of dCramp1 in suppressing transpositions is to avert transposon transcription. Supporting this hypothesis, single-molecule RNA fluorescent in situ hybridization (RNA-FISH) assay revealed strong activation of *HMS-Beagle* transcription in *dcramp1* depleted pupal ilea (fig. S5A). While *HMS-Beagle* started mobilizing in 5-day-old pupal ilea upon *dcramp1* depletion (Fig. 1B), we found that *HMS-Beagle* increased its transcription from pupa 3d (fig. S5A). These data indicate that, during somatic development, the timing of mRNA transcription and genomic integration of *HMS-Beagle* was completely different. Considering the most abundant transcripts of *HMS-Beagle* were detected at pupa 4d upon *dcramp1* depletion (fig. S5A), we extracted the total RNA at this time point and performed RNA-seq to quantify the expression of all transposons. Surprisingly, when comparing to control group, we found that more than half of the transposon families increased their transcription by more than 2-fold in *dcramp1* depleted ilea (fig. S5, B and C). Additionally, we found that transposon expression was positively corelated with novel insertions in *Drosophila* ileum (fig. S5D).

Previous studies have shown that retrotransposons can form double-stranded RNAs (dsRNAs) upon activation, which can be recognized by MDA5 receptor to induce immune responses(*44–46*). Leveraging strand-specific RNA-Seq method, we were aware of both sense and antisense transcripts produced by some retrotransposon families (fig. S5E), indicating their potential to form dsRNAs. The J2 antibody, which serves as a gold standard in dsRNA detection, showed strong immunostaining signals in *dcramp1* but not *white* depleted ilea (fig. S5F). As LTR-retrotransposons, upon activation, presumably they could generate linear double-stranded DNAs with an LTR at each end, requiring for transpositions(*47*). To examine whether the activated retrotransposons are able to generate dsDNAs, we applied END-seq, a sequencing assay capturing free dsDNA ends. END-seq showed strong signals at the ends of LTR-retrotransposons in *dcramp1* depleted hindguts (fig. S5G). Moreover, the density of END-seq is positively correlated with both expression and mobilizations of retrotransposons (fig. S5, H and I). With dsRNA/dsDNA produced and mobilizations generated upon transposon activation, *dcramp1* suppressed flies showed significantly higher lethality rates (fig. S5J). Given dCramp1 specifically suppresses transposon activity during ileum regeneration, the high lethality rate in *dcramp1* depleted flies was most likely caused by the defective regenerated hindguts.

### CRAMP1 suppresses transposon transcription in mouse fetal liver

Notably, when performing RNA-seq assay in *Cramp1*^-/-^ fetal livers, we discovered that many transposon families became highly active (fig. S3, E and F). When comparing two different time points, we found that much more transposons became transcriptionally active at E13.5d than at E11.5 in *Cramp1*^-/-^ fetal livers (fig. S3, E and F). At E13.5d, 36 transposons families increased their expression by more than 1.5-fold and 100 transposon families increased their expression by more than 1.2-fold but less than 1.5-fold (fig. S3F).

To further characterize the celluar and molecular alterations associated with the anemia phenotype in the absence of CRAMP1, we performed single cell RNA sequencing (scRNA-seq). Transcriptomic profiles were obtained from a total of 20,836 cells, of which 80% passed the quality control. Cells were clustered into 12 main cell types and annotated based on the expression of well-known marker genes (fig. S6, A and B)(*48*). After analyzing the cell type composition in mutant and control fetal livers, we observed a dramtic compositional perturbation of erythroid popuations in *Cramp1^-/-^* group (fig. S6C). Particularly, the erythroid progenitors showed 41.5% reduction (from 34.7% in *Cramp1*^+/-^ to 20.3% in *Cramp1*^-/-^), indicating an abnormal erythropoiesis in *Cramp1* mutated fetal liver (fig. S6, D and E). To examine whether the differed proportions are correlated with transposon activation, we also quantified the expression of transposable elements. The results revealed that many transposon families became highly active in the erythroid lineages, but not hepatocytes and endothelial cell, upon *Cramp1* mutation (fig. S6F). The scRNA-seq data and AZT treatment experiment suggested the potential effects of corrupted transposons on causing erythropoiesis defects.

### dCramp1/CRAMP1 promotes proper deposition of histone modifications to heterochromatin

Histone modifications, including both active and repressive ones, have long been considered as key factors to regulate transposon activity in somatic cells. Thus, we performed immunostaining assays to identify whether and which histone modifications could be affected by *dcramp1* depletion that leads to transposon activation in *Drosophila* ileum (Fig. 2A and fig. S7A). While majority of the histone modifications were not affected (fig. S7A), the localization of three repressive histone modifications marking heterochromatin is notably altered upon *dcramp1* depletion (Fig. 2A). In *dcramp1* depleted ilea, both H3K36me2 and H4K20me3 showed signal spreading from heterochromatin to euchromatin regions, broadly dispersing in the nucleus (Fig. 2A). More importantly, H3K9me3, the pivotal mark associated with constitutive heterochromatin, was also mis-localized in the nucleus upon *dcramp1* depletion (Fig. 2A). As reported previously, to establish heterochromatically repressed regions, H3K9me3 will subsequently recruit heterochromatin protein 1 (HP1) to these areas(*49–51*). Consistently, upon *dcramp1* depletion, the localization pattern of H3K9me3 and HP1 was completely different from control group, showing multiple foci in the nucleus (fig. S7B). Altogether, these data indicate that, during *Drosophila* hindgut regeneration, dCramp1 plays an important role in promoting heterochromatin formation for achieving transposon suppression.

**Fig. 2.**
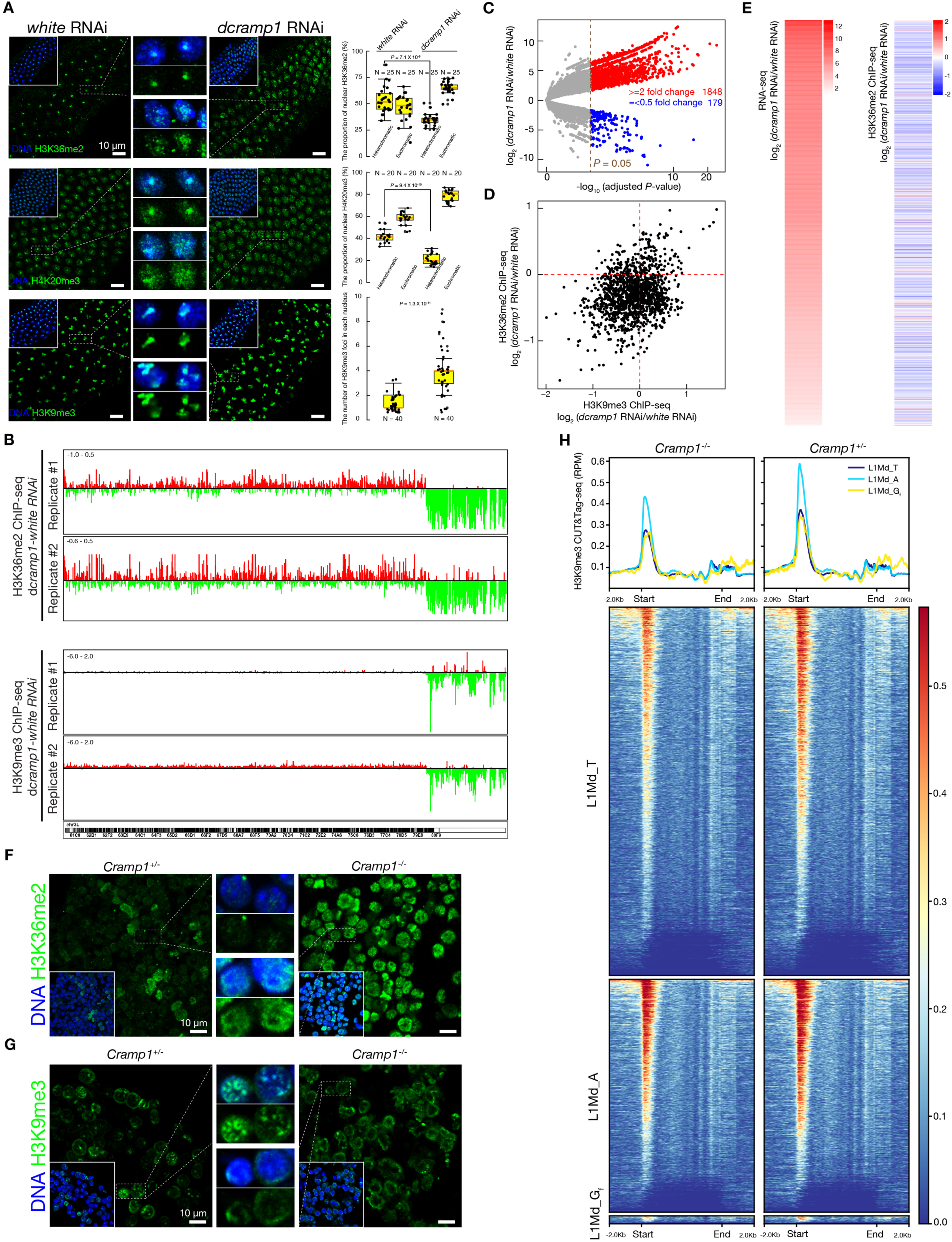
dCramp1/CRAMP1 promotes proper deposition of histone modifications to heterochromatin. A, Immunostaining to detect the nuclear signals of H3K36me2, H4K20me3 and H3K9me3 in *white* and *dcramp1* depleted pupal ilea. Box plots showing the relative fluorescence intensity of H3K36me2 and H4K20me3 and the numbers of H3K9me3 foci. The values of the fluorescence were measured by using ImageJ. The central lines represent median values, the box edges represent minimum and maximum values and the whiskers show the interquartile range of the data. Two-tailed *t*-tests were used to evaluate the difference between *white* and *dcramp1* depleted groups. Four-day-old pupae were used for immunostaining. B, Genome browser snapshots to depict H3K36me2 and H3K9me3 changes in *dcramp1* depleted 4-day-old pupal hindguts on chr3L, relative to *white* depleted pupal hindguts. Red bars above zero represent the gain of H3K36me2 and H3K9me3 signals. Green bars below zero represent the loss of H3K36me2 and H3K9me3 signals. C, Volcano plot showing differential transcription of locus-specific transposons between *white* and *dcramp1* depleted pupal hindguts. Red dots: Transposons that significantly increased their expression by more than 2 folds in *dcramp1* depleted hindguts. Blue dots: Transposon families that significantly reduced their expression by more than 2 folds in *dcramp1* depleted hindguts. D, Scatter plot showing the changes of H3K36me2 and H3K9me3 occupancy on activated transposons in *dcramp1* depleted 4-day-old pupal hindguts, relative to *white* depleted pupal hindguts. E, Heat map to depict the occupancy of H3K36me2 on activated transposons in *dcramp1* depleted 4-day-old pupal hindguts. These transposons showed 2-fold increased transcription upon *dcramp1* depletion. F and G, Immunostaining to detect the nuclear signals of H3K36me2 (F) and H3K9me3 (G) in *Cramp1*^+/-^ and *Cramp1*^-/-^ fetal liver at E13.5d. H, Metaplot and heat map for different transposon families of H3K9me3 CUT&Tag signal in reads per million (RPM) from *Cramp1*^+/-^ and *Cramp1*^-/-^ fetal TER119-positive cells.

Next, we performed chromatin immunoprecipitation followed by sequencing (ChIP-Seq) to determine the genome-wide profiling of H3K36me2 and H3K9me3. On the chromosomal level, in *dcramp1* depleted ilea, we observed that both H3K36me2 and H3K9me3 are dramatically depleted at pericentromeric regions (Fig. 2B and fig. S7, C and D). These pericentromeric regions in *Drosophila* genome are relatively gene poor and densely occupied with transposons and piRNA clusters (fig. S7C)(*52*). Next, we focused on analyzing these 1,848 transposons that significantly increased transcript abundance more than 2-fold in the genome upon *dcramp1* depletion (Fig. 2C). Among these activated transposons, 1,356 of them showed well-qualified H3K9me3 and H3K36me2 ChIP-seq signals (Fig. 2D). When intersecting the ChIP-seq data with transcriptionally activated transposons, we found that, 809 (59.7%) and 1,139 (84.0%) of transposons harbored decreased H3K9me3 and H3K36me2, respectively (Fig. 2, D and E and fig. S7E). These data reveal that, upon *dcramp1* depletion, H3K36me2 and H3K9me3 failed to properly deposit to transposon sequences, resulting in transposon derepression.

To illustrate whether CRAMP1 also regulates repressive histone modifications in fetal liver to control transposon activity, we performed the immunostaining for H4K20me3, H3K36me2 and H3K9me3. While no obvious differences were detected for H4K20me3, we observed strong increased signals of H3K36me2 in *Cramp1* mutated fetal liver (Fig. 2F and fig. S8A), consistent with the signals detected from *dcramp1* depleted *Drosophila* ileum (Fig. 2A). Additionally, much weaker H3K9me3 signals in heterochromatin were noticed in *Cramp1* mutated fetal liver, indicating heterochromatin formation was disrupted (Fig. 2G). Interestingly, we detected no differences of H3K36me2 in fetal kidney and heart between *Cramp1*^+/-^ and *Cramp1*^-/-^, suggesting the function of CRAMP1 in regulating repressive histone modifications is exclusive to fetal liver (fig. S8, B and C). Next, we sorted the TER119 positive cells (containing early erythrocytes based on scRNA-seq) from *Cramp1*^+/-^ and *Cramp1*^-/-^ E13.5 fetal liver and performed the CUT&tag experiments by H3K9me3 and H3K36me2 antibodies (fig. S8, D and E). We detected that the enrichment of H3K9me3 and H3K36me2 were clearly diminished for different transposon families in *Cramp1*^-/-^ (Fig. 2H and fig. S8F), especially the young LINE1 families (L1Md_T, L1Md_G_f_ and L1Md_A) that actively mobilized in fetal liver (Fig. 1H). To better understand the effects of reduced H3K9me3 and H3K36me2 deposition on increased transposon expression, we performed assay for transposase-accessible chromatin (ATAC) sequencing. We found LINE-1 retrotransposon showed more accessible at the promoter regions in *Cramp1*^-/-^ comparing to control group (fig. S8G).

Collectively, these results revealed that Cramp1 plays an evolutionally conserved role in establishing heterochromatic identities for transposon suppression.

### Cramp1 initiates linker histone H1 transcription by specifically binding to the sequence of histone gene cluster

Given the absence of dCramp1 in heterochromatin regions (fig. S1, B and C), we concluded that dCramp1 unlikely binds to transposon sequences to recruit the mediators of repressive histone modifications. One possibility is that Cramp1 regulates the gene expression regarding histone modifications. Indeed, we found that the expression of many histone genes was significantly decreased in *Cramp1* mutated fetuses (Fig. 3A and fig.S9A). Surprisingly, at E13.5d, among 44 genes whose expression reduced by more than 2-fold in *Cramp1* mutated fetal liver, 19 (43.2%) of them were histone genes (Fig. 3A). Notably, among those differentially expressed histone genes at E13.5d, we noticed that the top three decreased genes were coding linker histone H1, including *Hist1h1c*, *Hist1h1e* and *Hist1h1d* (Fig. 3A and fig.S9, A and B). Additionally, these three genes were the only reduced histone genes at E11.5d in *Cramp1* mutated fetal liver (Fig. 3A). As expected, the protein level of linker histone HIST1H1C, the top one decreased histone gene in *Cramp1* mutated fetal liver, was diminished (Fig. 3B). More precisely, *Hist1h1c*, *Hist1h1e* and *Hist1h1d* were significantly reduced in erythroid lineages as determined by scRNA-seq data (fig. S9, C and D). By comparing different fetal tissues, we discovered that *Hist1h1c*, *Hist1h1e* and *Hist1h1d* exhibited the most transcriptional level in liver (fig. S9E). Meanwhile, we found that these three genes showed the most significantly decrease in *Cramp1* mutated fetal liver (fig. S9F), which could be the reason why CRAMP1 mainly regulates histone modifications and suppresses transposon activity in fetal liver.

**Fig. 3.**
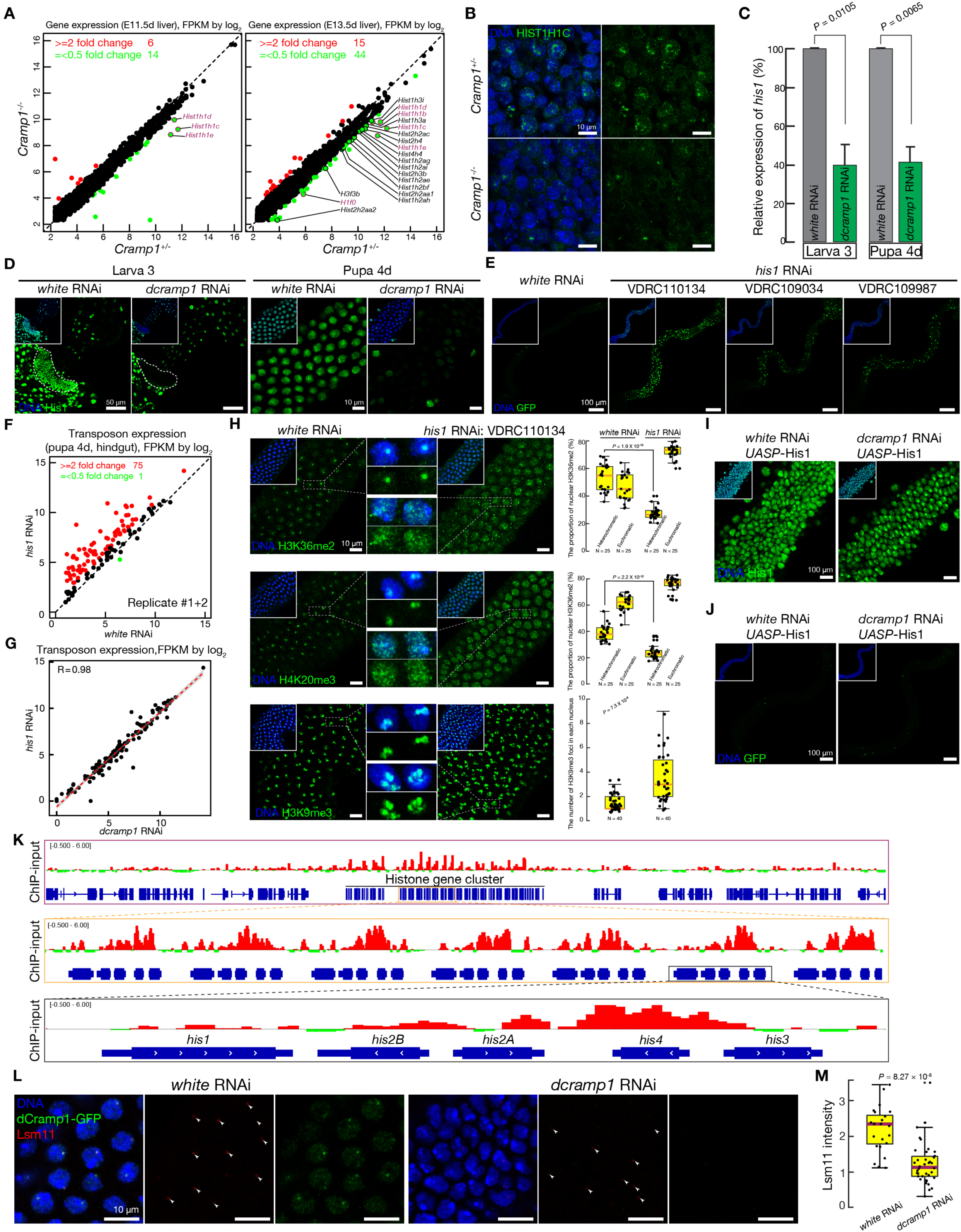
Cramp1 initiates linker histone H1 transcription by specifically binding to the sequence of the histone gene cluster. A, Transcriptome profiles from *Cramp1*^+/-^ and *Cramp1*^-/-^ fetal livers at E11.5d and E13.5d. Red dots: Genes that increased their expression by more than 2 folds in *Cramp1*^-/-^ fetal livers. Green dots: Genes that decreased their expression by more than 2 folds in *Cramp1*^-/-^ fetal livers. All of the histone genes are labeled and linker histone H1 genes are marked as purple. Data are merged from n = 2 biological replicates. B, Immunostaining to detect HIST1H1C production in *Cramp1*^+/-^ and *Cramp1*^-/-^ fetal livers at E13.5d. C, RT-qPCR to quantify *his1* expression in *white* and *dcramp1* depleted 4-day-old pupal hindguts based on the two-tailed *t*-test. Data are normalized to *rp49* (*RpL32*) expression; the bars report mean ± standard deviation for at least three biological replicates. D, Immunostaining to detect His1 production in larval and 4-day-old pupal hindguts upon *white* and *dcramp1* depletion. Dashed circles highlight the ring structure. E, Monitoring *HMS-Beagle* mobilization events from 1-day-old adults in *white* and *his1* depleted *Drosophila* ilea. Each GFP positive cell indicates *HMS-Beagle* mobilized at least once in the genome. Similar *HMS-Beagle* transpositions were detected from three different *his1* RNAi alleles. F, Scatter plot showing the transposon expression in *white* and *dcramp1* depleted 4-day-old pupal hindguts. Data are merged from n = 2 biological replicates. Red dots: Transposon families that increased their expression by more than 2 folds in *dcramp1* depleted hindguts. Green dots: Transposon families that reduced their expression by more than 2 folds in *dcramp1* depleted hindguts. G, Scatter plot showing the correlation of transposon transcription between *his1* and *dcramp1* depleted 4-day-old pupal hindguts. Pearson correlation coefficient is indicated. H, Immunostaining to detect the nuclear signals of H3K36me2, H4K20me3 and H3K9me3 in *white* and *his1* depleted 4-day-old pupal hindguts. Box plots showing the relative fluorescence intensity of H3K36me2 and H4K20me3 and the numbers of H3K9me3 foci. I, Immunostaining to detect His1 expression in *UASP*-His1 transgenic 4-day-old pupal hindguts upon *white* and *dcramp1* depletion. J, Monitoring *HMS-Beagle* mobilizations in *UASP*-His1 transgenic 4-day-old pupal hindguts upon *white* and *dcramp1* depletion. K, Genome browser snapshots to characterize the genomic binding loci of dCramp1 on chr2L from 4-day-old pupal hindguts. L, Immunostaining to detect dCramp1 and Lsm11 localization in *white* and *dcramp1* depleted 4-day-old pupal ilea. M, Box plots showing the fluorescence intensity of nuclear Lsm11 signals in *white* and *dcramp1* depleted 4-day-old pupal ilea. Note: H and M, the values of the fluorescence were measured by using ImageJ. The central lines of box plot represent median values, the box edges represent minimum and maximum values and the whiskers show the interquartile range of the data. Two-tailed *t*-tests were used to evaluate the difference between two groups.

Similarly, we discovered that, upon *dcramp1* depletion, only *his1* gene but not others was significantly decreased at both transcriptional and protein level in *Drosophila* hindguts (Fig. 3, C and D and fig. S10A). At the pupal stage, we noticed that His1 proteins are abundantly enriched at heterochromatin regions (Fig. 3D), suggesting a potential function of His1 in promoting heterochromatin formation. To test whether dCramp1 achieves transposon suppression through licensing linker histone H1 expression, we utilized three different RNAi alleles against *his1*, which could efficiently deplete *his1* expression in both pupal ileum and larval ring (fig. S10B). As expected, *his1* depletion led to not only *HMS-Beagle* mobilizations but also more than half of activated transposons in *Drosophila* ilea (Fig. 3, E and F and fig. S10, C and D). Notably, the aberrantly expressed transposons between *his1* and *dcramp1* depletion are highly corelated (Fig. 3G). Additionally, we found that *his1* suppressed *Drosophila* ilea generated more DNA breaks (fig. S10, E and F) and accumulated more dsRNAs in the nucleus (fig. S10G), most likely generated upon retrotransposon activation and mobilizations. Similar as *dcramp1* depleted ilea (Fig. 2A), we also discovered that *his1* depletion caused redistribution of H3K36me2 and H4K20me3 from repetitive regions and mis-localization of H3K9me3 in *Drosophila* ilea (Fig. 3H). More importantly, the activation of His1 in *dcramp1* depleted *Drosophila* ileum could fully silence *HMS-Beagle* activation and mobilizations (Fig. 3, I and J). These data indicate that His1 is the only downstream target for dCramp1 to execute the function in suppressing transposon mobility.

Given dCramp1 is a DNA binding protein, to address how dCramp1 promotes H1 expression, we dissected the hindguts from pupa 4d and performed dCramp1 ChIP sequencing. Surprisingly, we found that dCramp1 specifically binds the DNA sequences of histone gene cluster, with the highest enrichment at *his4* gene locus (Fig. 3K). As reported previously, the histone gene cluster is invariably colocalized with histone locus body (HLB), a non-membrane structure containing factors necessary for the transcription and processing of histone messenger RNAs(*53–55*). In the nucleus of both larval ring and pupal ileum, we noticed that dCramp1 is colocalized with Lsm11, a marker of the histone locus body (fig. S10, H and I). Moreover, we discovered that dCramp1 plays an essential role in nucleating histone locus body formation; upon *dcramp1* depletion, the accumulation of Lsm11 in histone locus body was significantly decreased (Fig. 3, L and M). Since the binding of dCramp1 to DNA is dependent on its SANT domain, deletion of this domain in dCramp1 led to the failure of DNA binding and subsequent degradation (fig. S10J). Collectively, our data indicated that dCramp1 stimulates linker histone H1 transcription by initiating histone locus body formation, resulting in transposon suppression.

### dCramp1 specifically executes its function during hindgut regeneration

Given *Drosophila* is a holometabolous insect, metamorphosis is initiated at the pupal stage, during which the larval ileum is first degenerated and then the adult one is reproduced(*34, 56*). Our data indicated that *dcramp1* RNAi efficiently depleted dCramp1 production in the anterior part of pylorus (ring) (Fig. 3D), the essential part for regenerating adult pylorus and ileum. Thus, we next tested when is the key time point that dCramp1 executes its function in promoting linker histone expression and transposon suppression (fig. S11A). Interestingly, when depleting *dcramp1* and *his1* specifically at the time windows before hindgut regeneration (larva, pupa 0d-2d), *HMS-Beagle* became active and mobilized in the ilea (fig. S11B). In contrast, only activating RNAi in fully regenerated ilea (pupa 3d-4d) led to similar *HMS-Beagle* transposition events among *white*, *his1* and *dcramp1* flies (fig. S11B). Additionally, specifically activating RNAi against *dcramp1* and *his1* at larval stage led to dispersing of H3K36me2 and H4K20me3, mis-localization of H3K9me3 and strong DNA breaks in regenerated ilea (fig. S11, C and D). Moreover, upon specifically depleting *dcramp1* at the larval stage, while the production of dCramp1 could be recovered well at 4-day-old pupal hindguts (fig. S11E), His1 expression was still diminished (fig. S11F). These results could not only explain why specifically activating *dcramp1* RNAi before regeneration induced transposon activation, but also indicate that ileum regeneration is a pivotal time window for dCramp1 mediated His1 transcription and transposon silencing.

### Linker histone H1 facilitates H3K36me2 mediated heterochromatin formation

To address which histone modification plays the core roles in initiating heterochromatin formation for achieving transposon silencing, we modeled the effects of histone modification mutations by inducing the production of H3.3K36M, H3.3K9I and H4K20M. While expressing H4K20M led to pupal lethality (data not shown), flies harboring activated H3.3K36M and H3.3K9I expression could develop to adult stage. As expected, H3.3K36M and H3.3K9I induction could efficiently abolish H3K36me2 and H3K9me3 in heterochromatin regions, respectively (fig. S12, A and B). We found that H3.3K9I activation did not affect the localization of H3K36me2 and H4K20me3, indicating normal establishment of heterochromatin in this scenario (Fig. 4A and fig. S12A). However, H3.3K36M production clearly interrupted the distribution of H4K20me3 and localization of H3K9me3 in chromatin (Fig. 4, A and B) and caused transcriptional activation of retrotransposons (fig. S12C). These results indicate that H3K36me2 is the most important histone modification to initiate heterochromatin formation for transposon silencing during *Drosophila* hindgut regeneration.

**Fig. 4.**
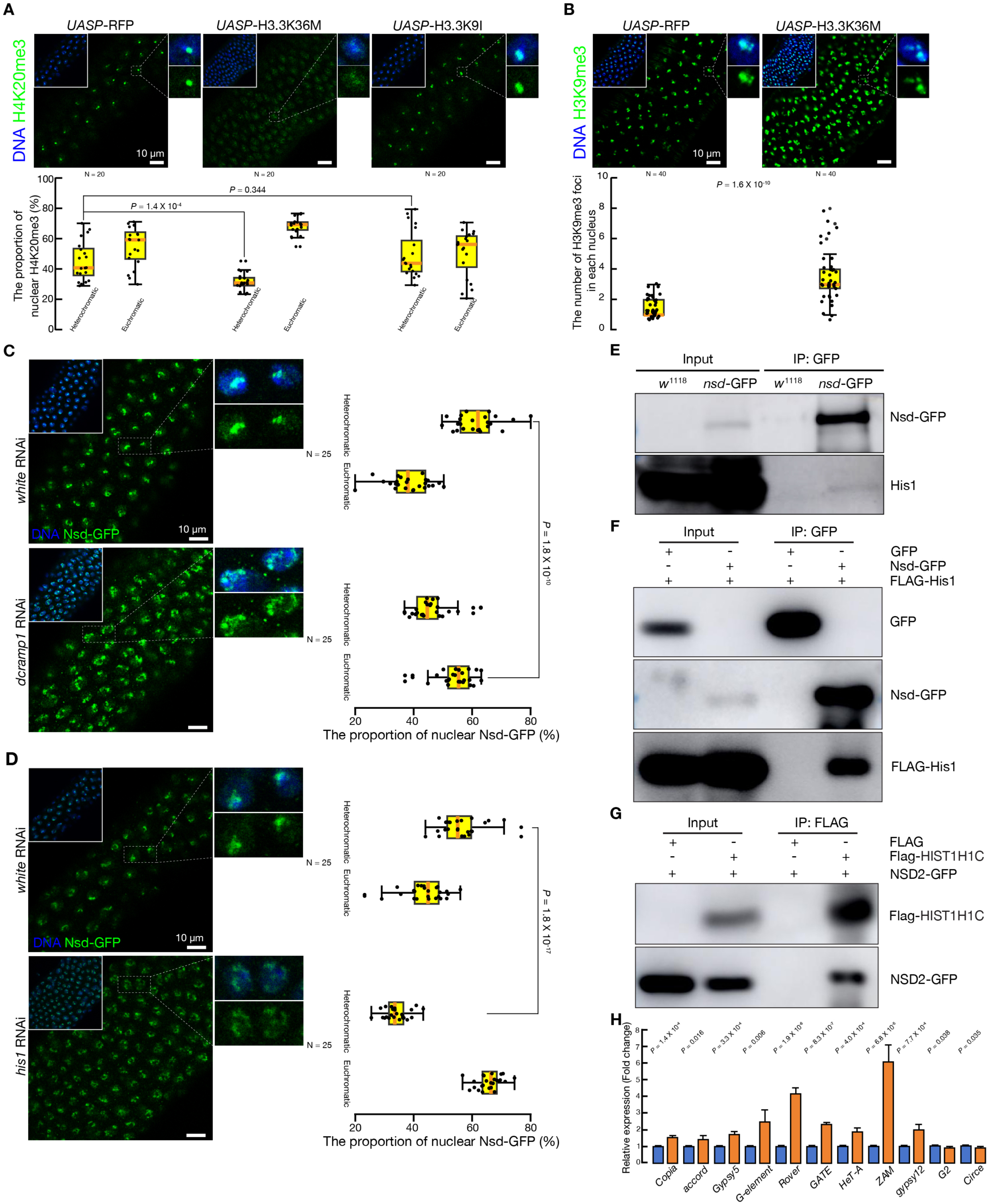
Linker histone H1 facilitates H3K36me2 mediated heterochromatin formation. A, Immunostaining to visualize nuclear signals of H4K20me3 in H3.3K36M and H3.3K9I 4-day-old pupal ilea. Box plots showing the relative fluorescence intensity of H4K20me3. *UASP*-RFP was used as control. RFP, H3.3K36M and H3.3K9I were driven by *byn*-Gal4. B, Immunostaining to visualize nuclear signals of H3K9me3 in H3.3K36M 4-day-old pupal ilea. Box plots showing the number of H3K9me3 foci. C and D, Immunostaining to detect the Nsd signals in *white*, *dcramp1* (C) and *his1* (D) depleted 4-day-old pupal ilea. Box plots showing the relative fluorescence intensity of Nsd. E and F, Co-immunoprecipitation to identify the interaction between *Drosophila* Nsd and His1. G, Co-immunoprecipitation to identify the interaction between mouse NSD2 and HIST1H1C. H, RT-qPCR to quantify the expression of retrotransposons in *white* and *nsd* depleted 4-day-old pupal hindguts based on the two-tailed *t*-test. Data are normalized to *rp49* (*RpL32*) expression; the bars report mean ± standard deviation for at least three biological replicates. Note: A, C and D, the values of the fluorescence were measured by using ImageJ. The central lines of box plot represent median values, the box edges represent minimum and maximum values and the whiskers show the interquartile range of the data. Two-tailed *t*-tests were used to evaluate the difference between two groups.

Two H3K36 methyltransferases, Nsd and Ash1, have been reported to catalyze the formation of H3K36me2(*23, 57*). To elucidate which methyltransferase contributes to H3K36me2 initiated heterochromatin formation during hindgut regeneration, we inserted a GFP tag into the N terminal of endogenous Nsd and Ash1. We observed that Ash1 was dispersed in the nucleus and Nsd was accumulated at heterochromatin regions, indicating Nsd is most likely the factor for H3K36 di-methylation during hindgut regeneration (Fig. 4, C and D and fig. S12D). Notably, both *dcramp1* and *his1* depletion disrupted the localization of Nsd at heterochromatin (Fig. 4, C and D). Given the linker histone H1 is also enriched at heterochromatin regions during hindgut regeneration, we hypothesized that the histone H1 might directly tether and recruit Nsd. Indeed, both the in vivo and in vitro Co-Immunoprecipitation (Co-IP) assay validated that the *Drosophila* Nsd could physically interact with His1 (Fig. 4, E and F, and fig. S12E). Accordingly, we found that mouse HIST1H1C, but not HIST1H1D and HIST1H1E, was also able to directly bind to NSD2, the homologue of *Drosophila* Nsd (Fig. 4G and fig. S12, F to H). Moreover, the transcription of many transposons became significantly activated in *nsd* depleted hindguts (Fig. 4H). Overall, our findings demonstrated that the linker histone H1 promotes H3K36me2 initiated heterochromatin formation by directly interacting with histone methyltransferase Nsd, eventually achieving transposon suppression.

## Discussion

Although the current view proposes that the transposon activity is suppressed by DNA methylation, repressive histone modifications and small RNA pathways, how transposon mobilizations are tightly controlled during somatic development remains largely unknown. By spatiotemporally tracking retrotransposon activity at the mobilization level during somatic development, we discovered that the small RNA pathways (from the results of RNAi screening, data not shown) contribute little, if any, to suppressing retrotransposon mobilizations. In contrast, we identified an evolutionally conserved mechanism in controlling transposon mobilizations during both *Drosophila* hindgut regeneration and fetal mouse erythropoiesis (fig. S13). Our data suggested that dCramp1 accumulates at the histone locus body to facilitate its formation and maintain its function, probably functioning as a “scaffolding” protein to promote linker histone H1 transcription.

Although transposon transcription and insertion have been used to assess transposon activity at different levels, little is known about how long it takes for retrotransposon to complete the transposition cycle. By monitoring *HMS-Beagle* transcription and mobilization, we found that, the timing of mRNA transcription and genomic integration was completely different. Our data indicated that, from transcribing the mRNA to integrating the genomic DNA, it took approximately two days for *HMS-Beagle* to accomplish the life cycle of transposition. Recently, Yang and his colleagues have discovered that alternative end-joining drives the extrachromosomal circular DNA (eccDNA) formation of LTR-retrotransposons, facilitating retrotransposon replication in *Drosophila* oocytes(*58*). During hindgut regeneration, within the two-day period, whether *HMS-Beagle* hijacks similar host machineries and prepares itself, such as eccDNA formation, to achieve mobilizations remains to be investigated.

By directly visualizing three repressive histone modifications during hindgut regeneration, we propose that heterochromatin is initiated by linker histone H1 mediated H3K36 di-methylation. Regarding this phenotype discovered during hindgut regeneration, less is known about how H3K36me2 regulates the deposition of H4K20me3 and H3K9me3 to repetitive sequences. One possible mechanism is that H3K36me2 might interact and recruit the histone methyltransferases to catalyze H4K20 and H3K9 trimethylation during hindgut regeneration. As we noticed similar results in fetal liver, future work will examine whether mammalian cells during fetal erythropoiesis employ a mechanism analogous to that of *Drosophila* hindgut regeneration to initiate heterochromatin formation. Although detailed mechanisms may differ, Cramp1 mediated linker histone H1 transcription, heterochromatin formation and transposon silencing during somatic development could potentially be a recurrent theme from flies to mammals.

## Method

### Fly strains and husbandry

Generally, all flies were maintained at 25 °C and grown on standard agar-corn medium. Given *HMS-Beagle* becomes most active at mobilization level at 18 °C, the flies used for RNAi screening and ubiquitously silencing *white*, *dcramp1* and *his1* were raised at 18 °C. To silence *white*, *dcramp1* and *his1* at the embryonic stage, eggs were laid and hatched at 29 ℃; larvae, pupae and adults were raised at 18 °C. To silence *white*, *dcramp1* and *his1* at the larval stage, eggs were laid and hatched at 18 ℃; larvae were raised at 29 ℃; pupae and adults were raised at 18 ℃. To silence *white*, *dcramp1* and *his1* at a specific time window of pupal stage, embryos and larvae were raised at 18 ℃; pupae with different developmental stages were placed at 29 ℃ for 24 hours and then transferred back to 18 ℃ until eclosed. Fly alleles for RNAi screening were bought from Vienna *Drosophila* Resource Center (VDRC), Bloomington *Drosophila* Stock Center (BDSC) and TsingHua Fly Center. We used *byn*-Gal4 to specifically deplete and activate gene expression in *Drosophila* hindguts.

### Mice

All mice experiments were performed in accordance with the guidelines of the Institutional Animal Care and Use Committee at the Shanghai Institute of Biochemistry and Cell Biology.

Mice were maintained in the specific pathogen-free grade and a temperature (23 ± 2 °C)- and light (12 hours light/dark cycle)-controlled animal facility at the Shanghai Institute of Biochemistry and Cell Biology. Mice were generated through standard mouse breeding procedures within the animal facility. Pregnant *Cramp1*^+/-^ female mice were killed at scheduled time points and fetal liver, kidney and heart tissues were collected for subsequent experiments.

### Cell culture and transfection

HEK293T cell line was purchased from the Core Facility for Stem Cell Research, CEMCS, CAS (CAT#: SCSP-502). HEK293T cells were cultured in DMEM (Thermo Fisher Scientific, CAT#: 11960500CP) supplemented with 10% FBS (Ausbian, CAT#: WS500T), 1% Glutamax (Thermo Fisher-Invitrogen, CAT#: 35050061), 1% Sodium Pyruvate 100mM Solution (Thermo Fisher-Invitrogen, CAT#: 11360070) and 1% Penicillin/Streptomycin (Thermo Fisher-Invitrogen, CAT#: 15140122). HEK293T cells were cultured at 37 °C in a humidified incubator with 5% CO_2_.

Transient transfection was performed by using Lipofectamine™ 3000 (Thermo Fisher-Invitrogen, CAT#: L3000008), following the manufacturer’s instruction. Plasmids and lipo3000 reagents were diluted in Opti-MEM (Thermo Fisher-Invitrogen, CAT#: 31985070) when transfecting the cells.

### Plasmid construction

The construct of *HMS-Beagle* transposition reporter (*HMS-Beagle*-TR) was made by Counter-Selection BAC Modification Kit (GENE BRIDGES, CAT#: K002). The BAC clone that contains the full length of *HMS-Beagle* is p[acman]-CH322-33A08, serving as the template to generate the GFP reporter. Full length of *HMS-Beagle* is 7059 bp, the sequence encoding Gag is from 1529 bp to 2927 bp, and the sequence encoding Pol is from 2867 bp to 6046 bp. The GFP reporter was inserted into *HMS-Beagle* between 6198 bp and 6199 bp, which will not disrupt the coding sequence.

The constructs of *dcramp1*-rescue (for making transgenic fly), *UASP*-*his1* (for making transgenic fly), *dcramp1*-GFP-WT (for making transgenic fly), *dcramp1*-GFP-SANT-KO (for making transgenic fly), *UASP*-H3.3K9I (for making transgenic fly), *UASP*-H3.3K36M (for making transgenic fly), *UASP*-H4K20M (for making transgenic fly), *nsd*-GFP (for making knock-in fly) and *ash1*-GFP (for making knock-in fly) were generated by Gibson Assembly and cloned into the EcoRI site of pCaSpeR3 vector.

To construct the plasmids for producing sgRNAs, the DNA fragments containing sgRNA sequences for generating knock-in flies were synthesized and cloned into pEASY-Blunt simple U6 vector.

The constructs of *nsd*-GFP (for in vitro transfection), *Nsd2*-GFP (for in vitro transfection), *his1*-FLAG (for in vitro transfection), *Hist1h1c*-FLAG (for in vitro transfection), *Hist1h1d*-FLAG (for in vitro transfection) and *Hist1h1e*-FLAG (for in vitro transfection) were generated by Gibson Assembly and cloned into the EcoRI and HindIII site of pUC57 vector.

All of the constructs were verified by colony PCR and Sanger sequencing. The plasmids were sent to the Core Facility of *Drosophila* Resource and Technology, CEMCS, CAS for generating transgenic or knock-in flies. The plasmids for generating transgenic flies were site-specifically landed into fly genome at attP2 and attP40 sites. *dcramp1*-GFP knock-in flies were generated by Qidong Fungene Biotechnology.

### Visualizing mobilization events from *HMS-Beagle* transposition reporter

To detect eGFP positive cells from *HMS-Beagle* mobilization reporter during somatic development, pupae and 1-2-day-old adult flies were heat-shocked at 37 ℃ for 45 minutes, then recovered at room temperature for 3-5 hours. *Drosophila* hindguts were dissected in cold PBS and fixed with 4% PFA at room temperature for 10 min. Fixed hindguts were then washed 3 times with PBST (PBS containing 0.1% Triton X-100), stained with DAPI and mounted with VECTASHIELD MOUNTING MEDIUM (VWR, CAT#: 101098-042). Given GFP possesses a nuclear localization signal, the cells shown GFP and DAPI co-localization were considered harboring real *HMS-Beagle* transposition events. Both pupal males and females were dissected and harbored similar number of GFP positive cells. Adult female flies were dissected to visualize GFP positive cells. Representative images were taken by the Leica TCS SP8 confocal microscope.

### Life span analysis

Newly eclosed flies (within 1-day-old) were collected from *white* and *dcramp1* depleted group. 10 males and 15 females were placed in each tube (two biologically independent experiments with 3 tubes in each experiment) and flipped to fresh food every other day, and the number of dead flies was counted when flipping the flies. All flies were raised at 25°C on regular medium with appropriate humidity. A comparison of the survival curves was completed in R 4.2.2 using the survival package (v3.5-7).

### RT-qPCR

The total RNA from *Drosophila* hindguts and mouse fetal tissues was extracted by using TRIzol Reagent (Thermo Fisher-Invitrogen, CAT#: 15596018) and precipitated by isopropanol. RNA was eluted in RNase-free water (Thermo Fisher Scientific, CAT#: 10977035) and the concentration was measured by NanoDrop. To digest any contaminated genomic DNA in RNA samples, 10 µg of total RNA was treated with 2 µl of Turbo DNase (Thermo Fisher Scientific, CAT#: AM2239) and incubated at 37 ℃ for 30 minutes. After Turbo DNase digestion, RNA was then purified by RNA Clean & Concentrator-5 (Zymo research, CAT#: R1016), and the concentration was measured by NanoDrop.

One µg of purified RNA was used for reverse transcription in 20 µl of reaction by using PrimeScript RT reagent Kit (TaKaRa, CAT#: RR047A). For each reaction, 1 µl of cDNA was used for qPCR by using HieffTM qPCR SYBR® Green Master Mix (Yeasen, CAT#: 11202ES03). qPCR reaction was performed by using Roche LightCycler 96 Instrument Real Time PCR System. *rp49* and *Actin* were used for normalizing *Drosophila* and mammal samples, respectively. *P*-values were calculated from at least three independent biological replicates using a two-tailed, paired *t*-test. The error bars on the graphs report the standard deviation for at least three independent biological replicates.

### RNA-seq library preparation

One µg of purified RNA was used for RNA-Seq library preparation by utilizing TruSeq Stranded Total RNA Library Prep kit (Illumina, CAT#: 20020596). Briefly, 1 µg of total RNA diluted to a final volume of 10 µl was used for library preparation. Add 5 µl of RBB (rRNA binding buffer) and 5 µl of RRM (rRNA removal mix) to RNA, mix well, then run the RNA denaturation program (68 °C, 5 minutes) on the thermal cycler. After denaturation, add 35 µl RRB (rRNA removal beads) to each sample to remove rRNA, then transfer the RNA to new tube. Add 99 µl RNAClean XP beads (Beckman Coulter, CAT#: A63987) to purify RNA and elute with 8.5 µl EPH (elute, prime, fragment high mix), then run the Elution 2-Frag-Prime program (94 °C, 9 minutes) on the thermal cycler for fragmenting the RNA. After fragmentation, add 7.2 µl FSA (First Strand Synthesis Act D Mix) and 0.8 µl SuperScript III (Thermo Fisher Scientific, CAT#: 18080044) to each sample, then place on the thermal cycler and run the Synthesize 1^st^ Strand Program (25°C, 10 minutes; 42°C, 15 minutes; 70°C, 15 minutes). After synthesizing the first strand cDNA, add 5 µl RBS (Resuspension buffer), 5 µl diluted CTE (end repair control, 1:50 dilution in RBS) and 20 µl SMM (second strand marking master mix) to synthesize the second strand cDNA (thermal cycler, 16 °C, 1 hour). Then, add 90 µl RNAClean XP beads to purify DNA, and elute with 17.5 µl RSB and transfer 15 µl of DNA to a new tube (−20 °C). Next Day, add 2.5 µl diluted CTA (A-tailing control, 1:100 diluted in RBS) and 12.5 µl ATL (A-tailing mix), mix well and place on the thermal cycler (37 °C, 30 minutes; 70 °C, 5 minutes) to adenylate 3’ end. After adenylation, add 2.5 µl diluted CTL (ligation control), 2.5 µl LIG (ligation mix) and 1 µl RNA adapters (10µM), then place on thermal cycler and run the LIG program (30 °C, 10 minutes). After ligation, add 5 µl STL (stop ligation buffer) to stop ligation, then purify DNA by using two rounds of RNAClean XP beads (round 1: 42 µl; round 2: 50 µl), elute the DNA with 52.5 µl and 22.5 µl of RSB, respectively. Transfer 19 µl eluted DNA to a new tube, add 5 µl PPC (PCR primer cocktail), 25 µl PMM (PCR master Mix) and 1 µl index, mix well and place on a thermal cycler and run the PCR program (98 °C 30 seconds; 15 cycles of 98 °C 10 seconds, 60 °C 30 seconds, and 72 °C 30 seconds; 72 °C 5 minutes) to amplify DNA fragments. Clean up enriched DNA fragments by using 50 µl RNAClean XP beads, elute with 20 µl RSB, and transfer 19 µl to a new tube. Send the libraries to Novogene and Sequanta for quality control and sequencing.

The expression of genes and transposons was analyzed and quantified using piPipes by family and SQuIRE (v 0.9.9.92) ‘Map’ and ‘Count’ function by locus. Sequencing reads were initially trimmed using cutadapt (v4.1) (DOI: https://doi.org/10.14806/ej.17.1.200) with parameters -e 0.1 -O 3 -m 55 --quality-cutoff 25, then the adapter-trimmed FASTQ files were used as input for both tools. SQuIRE calls STAR with options suited for TE analysis, quantification of reads per gene, and estimation of reads per TEs in a locus-specific manner. Differential expression analysis was done by DESeq2 using the raw count matrix of TEs from SQuIRE.

### Single cell RNA sequencing and analysis

Three whole fetal livers from *Cramp1*^+/-^ and *Cramp1*^-/-^ E13.5d embryos were dissected and stored in MACS Tissue Storage Solution (Miltenyi Biotec, CAT#: 130-100-008) provisionally. Livers were minced on ice and dissociated into single-cell suspensions by using dissociation enzyme (1×DPBS, 0.25% Trypsin (Hyclone, CAT#: SH30042.01) containing 10 µg/mL RNase-free DNase I (TaKaRa, CAT#: 2270B) and 5% FBS (Thermo Fisher Scientific, CAT#: SV30087.02). Livers were dissociated at 37 °C with a shaking speed of 50 rpm for about 40 minutes. Cell suspensions were filtered using a 40 µm nylon cell strainer and red blood cells were removed by 1x Red Blood Cell Lysis buffer (Thermo Fisher Scientific, CAT#: 00-4333-57). Dissociated cells were washed with 1×DPBS containing 2% FBS. Cells were stained with AO/Pl, and viability was assessed using a Countstar Fluorescence Cell Analyzer (Aber Instruments). Single cell RNA-seq library preparation and sequencing were performed by Benagen.

scRNA-seq data analysis: Sequenced reads were aligned to mouse reference genome mm10 by using CellRanger (v7.1.0) and then alignments lacking “CB” tags were filtered. scTE (v1.0) was employed to quantify the expression of genes and transposable elements. Expression matrices were loaded into Seurat R package (v4.4.0) and genes expressed in less than 3 cells were removed. Cells that did not meet the following criteria were excluded from the analysis: 1) The number of genes detected in each cell should be greater than 200 but less than 8000; 2) the UMIs of mitochondrial genes should be less than 20% of total UMI. RNA contaminations were removed via decontX (v0.99.3) and DoubletFinder (v2.0.3) was utilized to predict and eliminate doublets in the single cell data. A total of 16,629 cells and 32,145 features passed the filtration. The UMI counts of each cell were normalized using the “NormalizeData” function with a scale factor of 10,000 and the “FindVariableGenes” function was employed to identify variable genes. PCA was performed based on the variable genes and 30 PCs were loaded for UMAP dimensionality reduction. To find clusters, the same PCs were imported into “FindClusters” with resolution = 0.5. Next, the Wilcoxon Rank Sum test was performed to identify markers in each cluster. Cell types were manually assigned based on these markers with the help of CellMarker 2.0. To identify differentially expressed transposable elements (TEs) within each cluster, the “FoldChange” function was used. Furthermore, the expression levels of TEs were extracted from the data by using the “AverageExpression” function and the scCustomize (v1.1.3) was employed to allow for informative gene expression visualization.

### RNA FISH

Stellaris RNA FISH probe sets for *HMS-Beagle* is designed and purchased from LGC Bioresearch Technologies, and the probe of *HMS-Beagle* was used before. Briefly, 5-7 pupal hindguts were dissected in cold PBS and fixed with 4% formaldehyde for 10 minutes at room temperature. After fixation, hindguts were washed three times with PBS, and then immersed in 70% (v/v) ethanol for 8 hours at 4 °C. After that, hindguts were washed once with Wash Buffer A (LGC Biosearch Tech, CAT#: SMF-WA1-60) at room temperature for 5 minutes. Then samples were incubated with 50 µl Hybridization Buffer (LGC Biosearch Tech, CAT#: SMF-HB1-10) containing probe sets (125nM) for hybridization at 37 °C overnight. Next day, hindguts were washed twice with Wash Buffer A for 30 minutes at 37 °C and once with Wash Buffer B (LGC Biosearch Tech, CAT#: SMF-WB1-20) for 5 minutes at room temperature. Pupal hindguts were then mounted with VECTASHIELD MOUNTING MEDIUM (VWR, CAT#: 101098-042). Images were taken by the Leica TCS SP8 confocal microscope.

### Genomic DNA extraction

*Drosophila* tissues. 100 pairs of ovaries and 1,000 hindguts from *dcramp1* depleted females were dissected in cold PBS (ovaries and hindguts were stored at −80 °C until enough tissues were collected) and then homogenized in 400 µL Buffer A (100 mM Tris-HCl pH7.5, 100 mM EDTA, 100 mM NaCl, 0.5% SDS) with a disposable grinder until only cuticles remain. Add 1 µl of 10 mg/ml RNase A to samples and incubate at 37 °C for 10 minutes. Then, incubate the samples on a heat block at 65 °C for 30 minutes. After incubation, add 800 µL of Buffer B (28.6% 5 M potassium acetate and 71.4% 6 M lithium chloride) to each sample, mix well and place on ice for 1 hour. Afterwards, samples were centrifuged for 15 minutes at 12,000 rpm at room temperature. Transfer 1 ml of the supernatant into a new microcentrifuge tube. Add 600 µl of isopropanol to each sample, mix well and centrifuge at 12,000 rpm for 15 minutes at room temperature. Discard the supernatant, wash the pellet with 70% fresh-made ethanol, vortex and centrifuge at 12,000 rpm for 5 minutes. Discard the supernatant, air-dry, then elute the DNA with Nuclease-free water.

Mouse tails. Each sample was digested with 100 µl of lysis buffer (100 mM Tris-HCl pH7.5, 50 mM NaCl, 5 mM EDTA, 0.2% SDS) containing 1 µl of protease K (20 mg/ml, Thermo Fisher Scientific, CAT#: EO0491), incubating at 55 °C water bath overnight. Next day, DNA was precipitated by adding 300 µl of 100% ethanol, and washed once with 500 µl of 75% ethanol. Then, DNA was eluted with Nuclease-free water and measured by NanoDrop.

Fetal liver. E13.5d fetal livers from *Cramp1*^+/-^ and *Cramp1*^-/-^ embryos were dissected in ice-cold PBS, then livers were transferred to a sterile culture dish and washed once with 10 ml PBS on ice. Add 495 µl lysis buffer (100 mM Tris-HCl pH7.5, 50 mM NaCl, 5 mM EDTA, 0.2% SDS) and 5 µl Protease K to each sample, incubate at 55 °C water bath overnight. Next day, add 1 µl 10 mg/ml RNase A to each sample and incubate samples at 37 °C for 1 hour. Use Phenol/Chloroform/Isoamyl alcohol (25:24:1) to extract DNA and precipitate with ethanol. Elute DNA with 20 µl Nuclease-free water, and measure DNA concentration by Qubit dsDNA HS Assay Kit (Thermo Fisher Scientific, CAT#: Q32851).

### END-seq library preparation and data analysis

END-seq library preparation was performed as previously described. Briefly, around 150 4-day-old pupal hindguts were dissected in cold PBS within 1 hour. Then the hindguts were resuspended in cell suspension buffer and embedded in agarose plugs. Agarose plugs were treated with Proteinase K for 2 hours at 50 °C, then samples were incubated at 37 °C for 7 hours. After Proteinase K digestion, samples were treated with RNase A for 1 hour at 37 °C. The DNA within the plugs was blunted, A-tailed and ligated with a biotin-labelled hairpin adaptor containing a 3ʹ T overhang and Illumina’s p5 sequence. Subsequently, DNA was extracted from the melted agarose and sheared by sonication. Biotinylated DNA was enriched by streptavidin coated beads and the DNA fragments were ligated to a second adaptor containing Illumina’s p7 sequence followed by PCR amplification with Illumina sequencing primers for 22 cycles. Sequencing was performed on the DNBSEQ-T7 platform with 150-bp paired-end reads.

END-seq sequencing reads were firstly trimmed to remove adapters by cutadapt (v4.1) with parameters -e 0.1 -O 3 -m 55 --quality-cutoff 25, then reads were aligned to dm6 and TE sequences severally using bowtie (v1.2.1.1) with parameters -n 3 −l 50 -k 1. Functions ‘view’ and ‘sort’ of samtools (v.1.9) were used to convert and sort the aligned .sam files to sorted .bam files. After deduplication using Picard (v2.18.7) MarkDuplicates (https://broadinstitute.github.io/picard/), only the forward reads were retained using samtools ‘view’ function for subsequent analysis. .bam files were further converted to .bed files using the bedtools (v2.25.0) bamToBed command. BedGraph files were generated using bedtools genomecov, normalized by reads per million (RPM) and then converted into .bigWig files using bedGraphToBigWig from UCSC utilities.

### Immunostaining

*Drosophila* hindguts. Briefly, 5-7 pupal hindguts were dissected in cold PBS and fixed with 4% PFA in Buffer A (15 mM PIPES pH7.4, 80 mM KCl, 20 mM NaCl, 2 mM EDTA, 0.5 mM EGTA, 1 mM DTT, 0.5 mM Spermidine, 0.15 mM Spermine) for 10 minutes at room temperature. Hindguts were then washed twice with BAT Buffer (Buffer A, 0.1% TritonX-100). Hindguts were blocked with BAT buffer containing 10% Normal Donkey Serum (Absin, CAT#: abs935) at room temperature for 1 hour. Primary antibodies were diluted with NDS-BAT buffer and samples were incubated with primary antibodies at 4 °C overnight. Next day, hindguts were washed three times with BAT Buffer containing 2% BSA (Biosearch, CAT#: B-1000-100), 30 minutes each time. Then hindguts were incubated with secondary antibody (1:400) at 4 °C overnight. Next day, hindguts were washed once with BAT buffer for 30 minutes, once with BAT buffer containing DAPI for 30 minutes and once with BAT buffer for 30 minutes. Pupal hindguts were then mounted with VECTASHIELD MOUNTING MEDIUM and images were taken by the Leica TCS SP8 confocal microscope.

Mouse tissues. Fetal liver, heart and kidney from *Cramp1*^+/-^ and *Cramp1*^-/-^ E13.5d embryos were dissected and fixed with 4% PFA at 4 °C overnight. Next day, discard the fixation buffer, then wash mouse tissues with cold PBS twice. Discard PBS, add 70% ethanol and incubate for 1 hour at room temperature. Then successively add 75%, 80%, 95% and 100% ethanol to samples and incubate for 1.5 hours each time. After dehydration, transfer samples to preheating xylene, and incubate at 55 °C water bath for 1 hour. Replace the old xylene with new xylene and incubate the tissues for half an hour. Next, soak the mouse tissues in paraffin wax at 65 °C for 4 hours or overnight. Next day, embed tissues in liquid paraffin wax and cool to room temperature for sectioning. After sectioning, the samples were dewaxed in xylene twice, 20 minutes each time. Then rehydrate samples successively with 100%, 95%, 80%, 75% ethanol and water, for 5 minutes each time. After rehydration, place the samples in boiling sodium citrate antigen repair solution (10 mM sodium citrate, 0.05% Tween20 pH6.0) for 30 minutes, and cool naturally to room temperature. Then incubate samples in PBST (PBS, 0.5% Triton X-100) for 20 minutes to improve the penetration. Block the samples with PBST containing 10% NDS for 1 hour at room temperature. Primary antibodies were diluted with NDS-PBST and incubated with samples at 4 °C overnight. Next day, wash samples with PBST three times, 30 minutes each time. Dilute secondary antibodies (1:400) in NDS-PBST and incubate with samples for 1 hour at room temperature. Then the samples were washed three times with PBST and stained with DAPI. Finally, samples were mounted with VECTASHIELD MOUNTING MEDIUM and images were taken by the Leica TCS SP8 confocal microscope.

Antibodies used in immunostaining.

| ANTIBODY | SOURCE | IDENTIFIER | DILUTION |
| --- | --- | --- | --- |
| Anti-GFP | Abcam | CAT#: ab290 | 1:1000 (*Drosophila*) |
| Anti-GFP | Sigma-Aldrich | CAT#: AB16901 | 1:1000 (*Drosophila*) |
| Anti-H3K9me3 | Active motif | CAT#: ab39161 | 1:500 (*Drosophila*); 1:1000 (mouse) |
| Anti-H3K4me3 | Abcam | CAT#: ab8580 | 1:400 (*Drosophila*) |
| Anti-H3K9ac | Active motif | CAT#: ab39038 | 1:400 (*Drosophila*) |
| Anti-H3K9me2 | Active motif | CAT#: ab39683 | 1:400 (*Drosophila*) |
| Anti-H3K36me2 | Active motif | CAT#: ab39056 | 1:400 (*Drosophila*); 1:1000 (mouse) |
| Anti-H3K36me3 | Active motif | CAT#: ab61022 | 1:400 (*Drosophila*) |
| Anti-H4K20me3 | Active motif | CAT#: ab39671 | 1:400 (*Drosophila*, mouse) |
| Anti-H4K16ac | Abcam | CAT#: ab109463 | 1:400 (*Drosophila*) |
| Anti-H3K27me3 | Active motif | CAT#: ab39055 | 1:400 (*Drosophila*) |
| Anti-H3K27ac | Active motif | CAT#: ab39085 | 1:400 (*Drosophila*) |
| Anti-His1 | Active motif | CAT#: 61786 | 1:500 (*Drosophila*) |
| Anti-HIST1H1C | Abcam | CAT#: Ab17677 | 1:2000 (mouse) |
| Anti-dLsm11 |  | N/A | 1:200 (*Drosophila*) |
| Anti-HP1 | DSHB | CAT#: C1A9 | 1:1000 (*Drosophila*) |
| Anti-γH2Av | DSHB | CAT#: UNC93-5.2.1 | 1:400 (*Drosophila*) |
| Anti-γH2AX | CST | CAT#: 9718T | 1:1000 (mouse) |
| Anti-dsRNA | Jena Bioscience | CAT#: RNT-SCI-10010200 | 1:400 (*Drosophila*) |
| ALEXA FLUOR 488<br>DONKEY ANTI -Rb | Thermofisher-Invitrogen | CAT#: A21206 | 1:400 (*Drosophila*, mouse) |
| ALEXA FLUOR 488<br>DONKEY ANTI -Mo | Thermofisher-Invitrogen | CAT#: A21202 | 1:400 (*Drosophila*, mouse,) |
| ALEXA FLUOR 568<br>DONKEY ANTI-Rb | Thermofisher-Invitrogen | CAT#: A10042 | 1:400 (*Drosophila*, mouse) |
| ALEXA FLUOR 488 GOAT<br>ANTI -Chicken | Thermofisher-Invitrogen | CAT#: A11039 | 1:1000 (*Drosophila*, mouse) |

### TUNEL

The TUNEL experiments were performed by using DeadEnd Fluorometric TUNEL System (Promega, CAT#: G3250). Briefly, 5-7 pupal hindguts were dissected in cold PBS and fixed with 4% PFA for 10 minutes at room temperature. Wash hindguts in PBST (PBS, 0.1% Triton X-100) three times, 10 minutes each time. Discard PBST and resuspend the hindguts in 80 µl of Equilibration Buffer and incubate at room temperature for 10 minutes. Thaw the nucleotide mixture and rTdT on ice and prepare rTdT-Incubation-Buffer (90 µl Equilibration Buffer, 10 µl Nucleotide Mix, 2 µl rTdT Enzyme). Discard the Equilibration Buffer, then add the rTdT-Incubation-Buffer to each sample and incubate the samples at 37 °C for 3 hours, protecting samples from direct light exposure. Add 1 ml of 20 mM EDTA to terminate the reaction, then wash hindguts in 1 ml of PBST containing 5 mg/ml BSA for 10 minutes at room temperature. Discard supernatant, and mount pupal hindguts with VECTASHIELD MOUNTING MEDIUM. Images were taken by the Leica TCS SP8 confocal microscope.

### Fluorescence-Activated Cell Sorting (FACS)

Fetal livers from *Cramp1*^+/-^ and *Cramp1*^-/-^ at E13.5d were dissected in sterile culture dish with 1×DPBS solution. Livers were digest in following solution, 1×DPBS containing 0.25% Trypsin, 10 µg/ml DNase I (Thermo Fisher, CAT#: EN0523) and 5% Fetal Bovine Serum (Ausbian, CAT#: WS500T). Incubate and dissociate the livers at 37 ℃ with a shaking speed of 50 rpm for 40 minutes. Cell suspension was filtered using a 40 µm nylon cell strainer, then centrifuged at 500g for 5 minutes at 4℃. The cell pellets were re-suspended with 5 ml of pre-cooled 1×Red Blood Cell Lysis buffer (Thermo Fisher-Invitrogen, CAT#: 00-4333-57) and incubated on ice for 5 minutes. Dissociated cells were then washed with FACS buffer (2% FBS and 2mM EDTA in 1×DPBS) twice. Then, the cells were stained by Fixable Viability Dye eFlour506 (Thermo Fisher-Invitrogen, CAT#: 65-0866-14) and FITC anti-mouse TER-119 (Biolegend, CAT#: 116205) for 30 minutes at 4℃. Stained cells were washed twice with FACS buffer before cell sorting. Flow cytometry was performed using Sony MA900 Cell Sorter and the analysis was carried out with Flowjo (v10.8.1).

### ChIP-Sequencing library preparation and analysis

Dissect around 200 4-day-old *Drosophila* pupal hindguts in ice-cold PBS with needles, then transfer the hindguts to 500 µl cold PBS. Add 63.4 µl 16% PFA and incubate at room temperature for 10 minutes. After fixation, add 29.7 µl of 2.5 M glycine to a final concentration of 125 mM, and incubate at room temperature with rotation for 5 minutes. Hindguts were washed twice with ice-cold PBS, and once with Sonication Buffer (50 mM Tris-HCl pH8.0, 10 mM EDTA, 1% SDS, 1×Proteinase Inhibitor cocktail). Add 250 µl Sonication Buffer to each sample and sonicate samples for 4 rounds to shear DNA to an average fragment size of 200 - 1000 bp using Q-Sonic bioruptor (each round: duration time 15 minutes; on 20 seconds; off 40 seconds). Let the water in the Q-Sonic bioruptor cool to 4 °C after each round of sonication. After sonication, spin the samples at 16,000 g for 15 minutes at 4 °C, and transfer the supernatant to a new tube. Dilute the lysate with 9 volumes of 1×Dilution Buffer (16.7 mM Tris-HCl pH8.0, 1.1 mM EDTA, 1.1% Triton X-100, 1×Proteinase Inhibitor cocktail), and save 80 µl diluted lysate as Input.

Mix equal volumes of Dynabeads Protein A (Thermo Fisher-Invitrogen, CAT#: 10002D) and Protein G (Thermo Fisher-Invitrogen, CAT#: 10004D), and use 15 µl mixed beads for each immunoprecipitation. Wash beads twice with PBST (0.02% Tween20), and resuspend beads in 600 µl PBST. Add antibodies (anti-H3K9me3, Active motif, CAT#: ab39161; anti-H3K36me2 Active motif, CAT#: ab39056; anti-GFP, Abcam, CAT#: ab290) to each tube and incubate with beads at 4 °C with gentle rotation for at least 2 hours. After incubation, wash beads twice with PBST, then add diluted lysate, and incubate at 4 °C with gentle rotation overnight.

Next day, wash the samples twice with Wash Buffer A (20 mM Tris HCl pH8.0, 2 mM EDTA, 150 mM NaCl, 0.1% SDS, 1% Triton X-100), twice with ice-cold Wash Buffer B (20 mM Tris HCl pH8.0, 2 mM EDTA, 500 mM NaCl, 0.1% SDS, 1% Triton X-100), Wash Buffer C (10 mM Tris HCl pH8.0, 1 mM EDTA, 0.25 M LiCl, 1% Sodium deoxycholate, 1% NP-40) and TE Buffer (10 mM Tris-HCl pH8.0, 1 mM EDTA) sequentially. Add 80 µl Direct Elution Buffer (10 mM Tris-HCl pH8.0, 300 mM NaCl, 5 mM EDTA, 0.5% SDS) to each IP sample, and 3 µl of 5 M NaCl into saved Input samples. Incubate the IP and Input samples at 65 °C for 6 hours or overnight for de-crosslinking. Then add 1 µl of 10 mg/ml RNase A (Thermo Fisher Scientific, CAT#: EN0531) to each sample and incubate for 30 minutes at 37 °C. Add 2.5 µl of 20 mg/ml Proteinase K to each tube and incubate for 2 hours at 55 °C. Purify the DNA by using Phenol/Chloroform and precipitate DNA by using ethanol. Elute DNA in 10 µl water and measure DNA concentration by Qubit dsDNA HS Assay Kit.

Prepare the libraries for ChIP-Seq by using TruePrep DNA Library Prep Kit V2 for Illumina® (Vazyme, CAT#: TD502/TD503). Briefly, 5 ng/1 ng of purified DNA diluted to a final volume of 11 µl was used for DNA library preparation. Add 4 µl of TTBL (TruePrep Tagment Buffer L) and 5 µl of TTE (TruePrep Tagment Enzyme) Mix V5/V1 to each sample, mix well and incubate the samples at 55 °C for 10 minutes. After incubation, add 5 µl of TS (Terminate Solution) buffer and place the samples at room temperature for 5 minutes to stop the reaction. Finally amplify the DNA fragments by adding TAB (TruePrep Amplify Buffer), TAE (TruePrep Amplify Enzyme) and indexes. The libraries were purified by RNAClean XP beads and sent to Novogene for quality control and sequencing.

ChIP-seq data analysis. Adapters were trimmed from ChIP-seq reads via cutadapt with parameters -e 0.1 -O 3 -m 55 --quality-cutoff 25, and then using bowtie2 (v2.3.1) with default parameters aligned reads to dm6 and TE sequences. Keep high quality alignments into sorted.bam files via samtools ‘view’ function with parameters -h -b -f 3 -F 12 -q 20 -F 256 and ‘sort’ function. Duplications were removed using Picard MarkDuplicates and the .bam files were further converted to .bed files using the bedtools bamToBed command. BedGraph files were generated using bedtools genomecov, normalized by reads per million (RPM) and then converted into .bigWig files using bedGraphToBigWig from UCSC utilities.

### ATAC-seq library preparation and data analysis

One hundred thousand cells were collected through Flow cytometry for ATAC-seq library preparation. These libraries were performed by using Hyperactive ATAC-Seq Library Prep Kit for Illumina (Vazyme, CAT#:TD711). Briefly, cells were washed by TW Buffer (Tagment DNA & Wash Buffer) twice and then resuspended in Lysis Buffer (Resuspension Buffer, 10% NP40, 10% Tween20 and 1% Digitonin). Add transposition reaction mix (TW Buffer, 10% Tween20, 1% Digitonin, 5 × TTBL (TruePrep Tagment Buffer L), TTE (TruePrep Tagment Enzyme) Mix V50 and Nuclease-free water) to the samples and incubate the samples at 37℃ for 30 minutes, then stop the reaction by adding Stop Buffer. After that, the DNA fragments were purified by using ATAC DNA Extract Beads. Add the Amplification Mix (2×CAM, Illumina Primer 1, Illumina Primer 2) to DNA sample and amplify the libraries on thermocycler. The libraries were purified by RNAClean XP beads and sent to Novogene for quality control and sequencing.

The ATAC-seq sequencing reads were pre-processed using cutadapt with parameters -e 0.1 -O 3 -m 55 --quality-cutoff 25 to filter out adapter sequences. The reads were then mapped to the mm10 reference genome using bowtie2 with parameters --very-sensitive -X 2000. Reads mapping to the mitochondrial chromosome were filtered and duplicate reads were removed using Picard MarkDuplicates. Functions ‘view’ and ‘sort’ of samtools were used to convert and sort the aligned .sam files to sorted .bam files. .bam files were further converted to .bed files using the bedtools bamToBed command. All reads aligning to the “+” strand were offset by +4 bp, and all reads aligning to the “-” strand were offset −5 bp. BedGraph files were generated using bedtools genomecov, normalized by reads per million (RPM) and then converted into .bigWig files using bedGraphToBigWig.

### CUT&Tag library preparation and data analysis

One hundred thousand cells were collected through Flow cytometry for ATAC-seq library preparation. These libraries were prepared by using Hyperactive Universal CUT&Tag Assay Kit for Illumina Pro (Vazyme, CAT#:TD904). Briefly, cells were washed and resuspended by Wash Buffer. Cells were then incubated with ConA Beads Pro (Concanavalin A-coated Magnetic Beads Pro) for 10 minutes at room temperature. Add the primary antibodies (anti-H3K9me3, Active motif, CAT#: ab39161 and anti-H3K36me2, Active motif, CAT#: ab39056) to the samples and incubate at 4 ℃ overnight. Next day, dilute the secondary antibodies (Goat Anti-Rabbit IgG H&L) with the ratio of 1:100, then add the secondary antibodies into the samples and incubate for 1 hour at room temperature. After incubation, wash the beads with Dig-wash Buffer (Wash Buffer with 5% Digitonin) twice and then incubate the samples with pA/G-Tnp Pro (Hyperactive pA/G-Transposon Pro) for 1 hour at room temperature. After that, add TTBL (TruePrep Tagment Buffer L) to the samples and incubate at 37 ℃ for 60 minutes, then add 10% SDS into the samples to stop the tagmentation reaction. DNA fragments were purified by DNA Extract Beads Pro and the libraries were amplified by using Amplification Mix (2×CAM, Illumina Primer 1, Illumina Primer 2). The final libraries were purified by RNAClean XP beads and sent to Novogene for quality control and sequencing.

We used cutadapt with parameters -e 0.1 -O 3 -m 55 --quality-cutoff 25 to trim adapters, and then using bowtie2 with default parameters aligned reads to mm10 reference genome. Keeping properly and primarily mapped reads into sorted.bam files via samtools ‘view’ function with parameters -h -b -f 3 -F 12 -F 256 and ‘sort’ function. Duplications were removed using Picard MarkDuplicates and the .bam files were further converted to .bed files using the bedtools bamToBed command with blacklisted regions filtered. BedGraph files were generated using bedtools genomecov, normalized by reads per million (RPM) and then converted into .bigWig files using bedGraphToBigWig.

### Heatmaps and average profile plots

Coordinates of repetitive elements were extracted from mm10 annotations downloaded form RepeatMasker and filtered based on length to obtain a reasonable number of elements (LINE and LTR: greater than 5kb; DNA: greater than 500bp; SINE: greater than 300bp).

We utilized deepTools (v 3.2.1) computeMatrix to compute normalized read coverage across these repeat elements. The central regions were length-normalized to 5 kb with flanking regions extending 2 kb from the start and end positions. Subsequently, heatmaps were generated using deepTools plotHeatmap and the rows in the heatmap were sorted in descending order based on the total signal intensity.

### Co-Immunoprecipitation

The plasmids producing proper proteins were transfected to HEK293T cells by Lipo3000. Forty-eight hours post-transfection, cells were digested with trypsin and harvested. Cells were washed once with ice-cold PBS. Discard the PBS and add 600 µl lysis buffer (50 mM Tris-HCl pH7.4, 150 mM NaCl, 0.5% Triton X-100, 1×Protease Inhibitor cocktail, 1 mM PMSF), placing the samples on ice for 30 minutes. After lysing, spin the samples at 16,000 g for 15 minutes at 4 °C, and transfer the supernatant to a new tube, saving 5% of the lysate as Input.

Zero-12h of NSD-GFP knock-in and *w*^1118^ *Drosophila* embryos were collected for Co-IP assay. Embryos were washed twice with PBS. Add 600 µl lysis buffer (50 mM Tris-HCl pH7.4, 150 mM NaCl, 0.5% Triton X-100, 1×Protease Inhibitor cocktail, 1 mM PMSF), then place the samples on ice for 30 minutes. After lysing, spin the samples at 16,000 g for 15 minutes at 4 °C, and transfer the supernatant to a new tube, saving 5% of the lysate as Input.

Add 6×SDS protein loading buffer to Input and boil for 10 minutes, then store at −20 °C. The lysate was then transferred to the tubes containing 10 µl Sepharose anti-GFP antibody (Abcam, CAT#: ab69314) or 1 µl anti-FLAG antibody (Abcam, CAT#: ab236777) which is already incubated with 15μL Protein A/G beads at 4 °C for 3 hours. Incubate samples at 4 °C with gentle rotation overnight. Next day, wash IP samples five times with ice-cold cell lysis buffer and elute them with 2×SDS protein loading buffer, boil the IP samples at 95 °C for 10 minutes.

Run IP and Input samples on 8% SDS-PAGE gels, then transfer the proteins to 0.45 µm nitrocellulose membrane (Cytiva, CAT#: 10600002). After transferring the proteins, add 5% milk and incubate the samples at room temperature for 1 hour to block the membrane. Dilute primary antibodies with Dilution Buffer (Epizyme, CAT#: PS114) and incubate the samples at 4 °C overnight. Next day, wash the membrane with TBST three times, 10 minutes each time. Dilute secondary antibodies (1:5000) in 5% milk and incubate the samples at room temperature for 1 hour. Wash the membrane with TBST three times, 10 minutes each time. Finally, proteins were stained by using Tanon High-sig ECL Western Blotting Substrate (ABclonal, CAT#: 180-5001). Antibodies used in Co-immunoprecipitation and western blot.

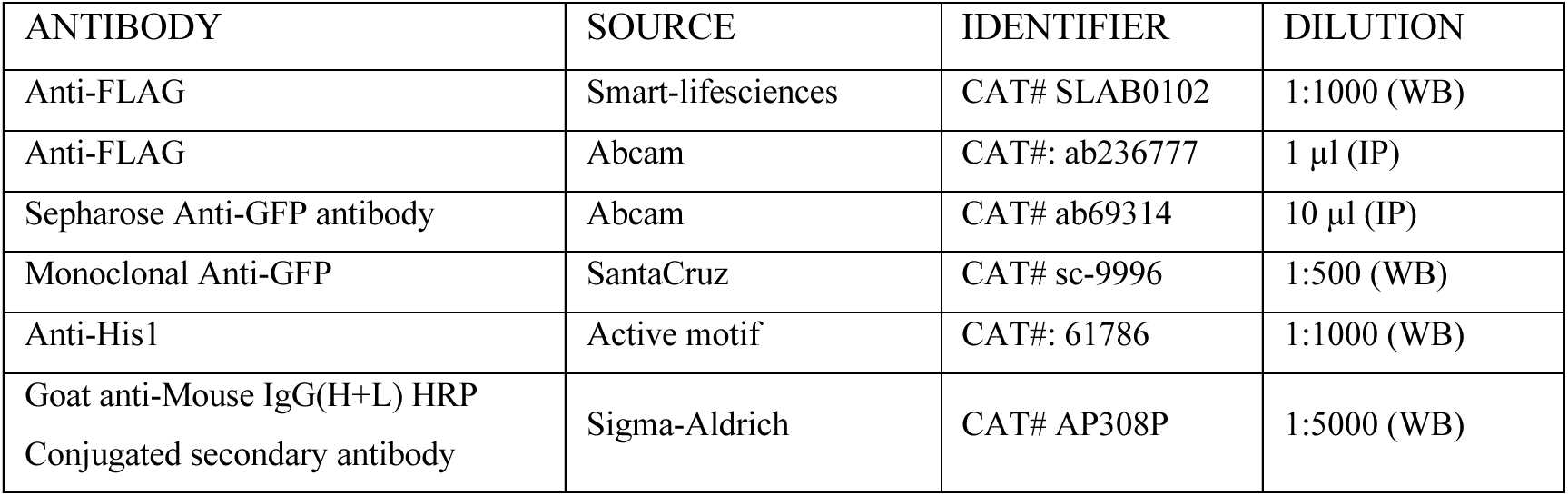

### AZT treatment

Dilute 3′-Azido-3′-deoxythymidine (AZT, Sigma Aldrich, CAT#: A2169) to 5 mg/mL in Nuclease-free water and filter by 0.22 μm Syringe Driven Filters (Jet Bio-Filtration, CAT#: STEM-GC-3374-Y). Aliquot AZT and store them at −20 °C. 50 mg/kg/day AZT was administered daily by gavage to around 10-week-old pregnant females from E7.5d to E14.5d, then no further treatment was performed. At E15.5d, the pregnant females were dissected and embryos were collected for measuring the weights and dissecting the fetal livers.

